# A systemic approach provides insights into the salt stress adaptation mechanisms of contrasting bread wheat genotypes

**DOI:** 10.1101/741090

**Authors:** Diana Duarte-Delgado, Said Dadshani, Heiko Schoof, Benedict C. Oyiga, Michael Schneider, Boby Mathew, Jens Léon, Agim Ballvora

## Abstract

Bread wheat is one of the most important crops for human diet but the increasing soil salinization is causing yield reductions worldwide. Physiological, genetic, transcriptomics and bioinformatics analyses were integrated to study the salt stress adaptation response in bread wheat. A comparative analysis to uncover the dynamic transcriptomic response of contrasting genotypes from two wheat populations was performed at both osmotic and ionic phases in time points defined by physiologic measurements. The differential stress effect on the expression of photosynthesis, calcium binding and oxidative stress response genes in the contrasting genotypes supported the greater photosynthesis inhibition observed in the susceptible genotype at the osmotic phase. At the ionic phase genes involved in metal ion binding and transporter activity were up-regulated and down-regulated in the tolerant and susceptible genotypes, respectively. The stress effect on mechanisms related with protein synthesis and breakdown was identified at both stress phases. Based on the linkage disequilibrium blocks it was possible to select salt-responsive genes as potential components operating in the salt stress response pathways leading to salt stress resilience specific traits. Therefore, the implementation of a systemic approach provided insights into the adaptation response mechanisms of contrasting bread wheat genotypes at both salt stress phases.

**Highlight:** The implementation of a systemic approach provided insights into salt stress adaptation response mechanisms of contrasting bread wheat genotypes from two mapping populations at both osmotic and ionic phases.

## Introduction

Bread wheat (*Triticum aestivum* L.) is a key staple crop for global food security and to feed the world population by 2050 its production needs to be increased substantially (Curtis and Halford, 2014). Therefore, breeding programs should emphasize on the genetic improvement of complex traits to increase yield potential under growth limiting conditions (Hawkesford *et al*., 2013). The genetic studies of complex traits in wheat are challenging because it is an allohexapolyploid species containing three sub genomes (AABBDD) with highly repetitive DNA sequences (85%) and a total genome size of 16 Gb (Loginova and Silkova, 2018; IWGSC, 2018). Efforts from the IWGSC (International Wheat Genome Sequencing Consortium) have resulted in the release of an annotated and highly contiguous chromosome-level assembly sequence draft of the Chinese Spring cultivar representing 94% of the whole genome (Shi and Ling, 2018; IWGSC; Borrill *et al*., 2019).

The world is losing 2000 hectares of arable soil daily due to salt-induced degradation which is a serious threat for global food security (Zaman *et al*., 2016). Among all the abiotic stress factors, soil salinity can cause significant yield reductions and decreased grain quality in wheat (Zheng *et al*., 2009). The salt stress adaptation response is a complex trait since the changes caused in key physiological processes are the effect of a coordinated action of gene networks in several metabolic pathways (Tuteja, 2007; Gupta and Huang, 2014). Therefore, the development of cultivars with increased salinity tolerance can be facilitated by a better understanding of the mechanistic basis underlying the stress adaptation response. This objective can be achieved through the integration of a range of experimental and statistical methods from various disciplines such as genetics, transcriptomics, bioinformatics and stress physiology to develop systemic approaches to gain a comprehensive understanding of relevant agronomic complex characters as the tolerance to saline conditions (Civelek and Lusis, 2014).

High salinity leads to physiological drought conditions, causes ion toxicity and cell oxidative damage that affect the plant growth (Tuteja, 2007; Gupta and Huang, 2014). Thus, plant growth response to salinity comprises two phases. The first corresponds to the osmotic phase which is independent of the sodium accumulation in tissues. The rapid and often transient impact on plant growth in this phase is attributed to the osmotic effect of the salts in the rhizosphere because of a reduced water potential (Ismail *et al*., 2014; Parihar *et al*., 2015; Julkowska and Testerink, 2015). Consequently, the osmotic tolerant plants have the ability to adapt to the drought aspect of the stress through the maintenance of the stomatal conductance and the leaf turgor (Carillo *et al*., 2011). The early signaling events in the osmotic phase that occur within seconds to hours of salt stress exposure are crucial for the acclimation response of the plants (Julkowska and Testerink, 2015). A model proposes that in the osmotic phase the root senses salt stress and second messengers as ROS (Reactive Oxygen Species) and Ca^2+^ are spread as signals to the aerial parts to trigger adaptive responses to cope with the Na^+^ ions that reach photosynthetic tissues and cause toxic effects in the following stress phase (Köster *et al*., 2018). Second, the ionic phase continues as a result of salt accumulation in leaves and takes days or even weeks to be manifested. In this phase the senescence of older leaves is caused by the inability of the plant to tolerate the toxic concentrations of salts in the tissues (Roy *et al*., 2014; Ismail *et al*., 2014). To reduce the toxicity effects, tolerant plants can excrete Na^+^ from leaves through the roots or compartmentalize Na^+^ and Cl^−^ in vacuoles to avoid toxic concentrations in the cytoplasm (Munns and Tester, 2008; James *et al*., 2012). Hence, the limiting effect of salinity on crop productivity in both stress phases is mainly due to its effect on the photosynthetic process which results in a substantial decrease in the biomass accumulation (Demetriou *et al*., 2007; Silveira and Carvalho, 2016).

The genetic mapping studies have allowed the detection of statistical associations of molecular markers with phenotypic values of a quantitative trait to find locations of quantitative trait loci (QTL) in the genome of a given species (Kang *et al*., 2016; Ishikawa, 2017). Several mapping analyses in bread wheat have identified QTL with effect on salt stress-related traits (Xu *et al*., 2013; Hussain *et al*., 2017; Dadshani, 2018; Oyiga *et al*., 2018). The recent studies by Oyiga *et al*. (2018) and Dadshani (2018) identified QTL for salt stress-related traits at germination, seedling and adult stages in association and advanced backcross-QTL (AB-QTL) mapping populations, respectively. Because of the limited mapping resolution of these studies, the linkage disequilibrium (LD) blocks where the QTL are localized contain many genes that are potential causal quantitative trait genes (QTGs) influencing the trait variation. A combination of genetic mapping analyses with transcriptomics can greatly reduce the number of these genes and provide with strong candidate QTGs for the identified QTL (González-Prendes *et al*., 2017; Ishikawa, 2017). In addition, transcriptomics analyses allow to identify the pathways that are regulated in the osmotic and ionic phases and give insights into the salt stress regulation of the photosynthesis from contrasting genotypes (Chaves *et al*., 2009; Liu *et al*., 2019). A time course transcriptomic analysis is therefore appropriate to identify key events during the salt stress response (Zhang *et al*., 2016; Liu *et al*., 2019).

The Next-Generation Sequencing platforms comprise rapidly evolving high throughput methods which are becoming the election to provide new insights into the whole genome, transcriptome, and epigenome of plants to assist the breeders to understand the biological function of the genes (Yadav *et al*., 2018). In contrast to regular RNA-seq, the Massive Analysis of cDNA 3’-ends (MACE) sequencing protocol is an alternative to perform transcriptomic analyses where a single sequence fragment represents one transcript (Hrdlickova *et al*., 2017; Tzfadia *et al*., 2018). Therefore, the output from this protocol is digital, strand-specific, and the quantification of the expression is simpler and more precise in each library (Asmann *et al*., 2009; Hrdlickova *et al*., 2017). This sequencing strategy can also contribute to better define gene models towards the 3’-ends (Tzfadia *et al*., 2018).

This study presents a systemic approach that integrates the dynamic transcriptomic responses of salt-tolerant and salt-sensitive wheat genotypes with photosynthetic activity and genetic mapping studies. The main goal of our study was to provide insights into the molecular mechanisms of the salt stress response in bread wheat through a comparative transcriptomic analysis and the identification of potential candidate genes operating in salt stress response pathways.

## Methods

### Contrasting genotypes from the mapping populations and tissue sampling

The elite German winter wheat cultivar Zentos (salt-tolerant; Belderok *et al*., 2000) and the synthetic genotype Syn86 (salt-susceptible; Lange and Jochemsen, 1992) were the contrasting parents used to study the foliar transcriptome during the osmotic stress response (Kunert *et al*., 2007; Dadshani, 2018). Altay2000 (salt-tolerant; Altay, 2012) and Bobur (salt-susceptible; Amanov, 2017; available at http://www.iwwip.org/images/dosya/636220600428830708.pdf) were the contrasting Turkish and Uzbeq winter wheat cultivars used to study the transcriptome during the ionic stress response, respectively (Oyiga *et al*., 2018). A differential photosynthesis response of Syn86 and Zentos was revealed by the sensor-based measurement of the time course of the photosynthesis rate performed under hydroponics conditions with 100 mol m^−3^ NaCl (Dadshani, 2018). This study allowed the identification of the “turning points” of the photosynthesis rate from 0 to 45 min after stress exposure (ASE). As “turning points” were considered the time points with maximum variation response, as revealed by the change of direction from the curve slope (Fig. 1A).

**Fig. 1.**
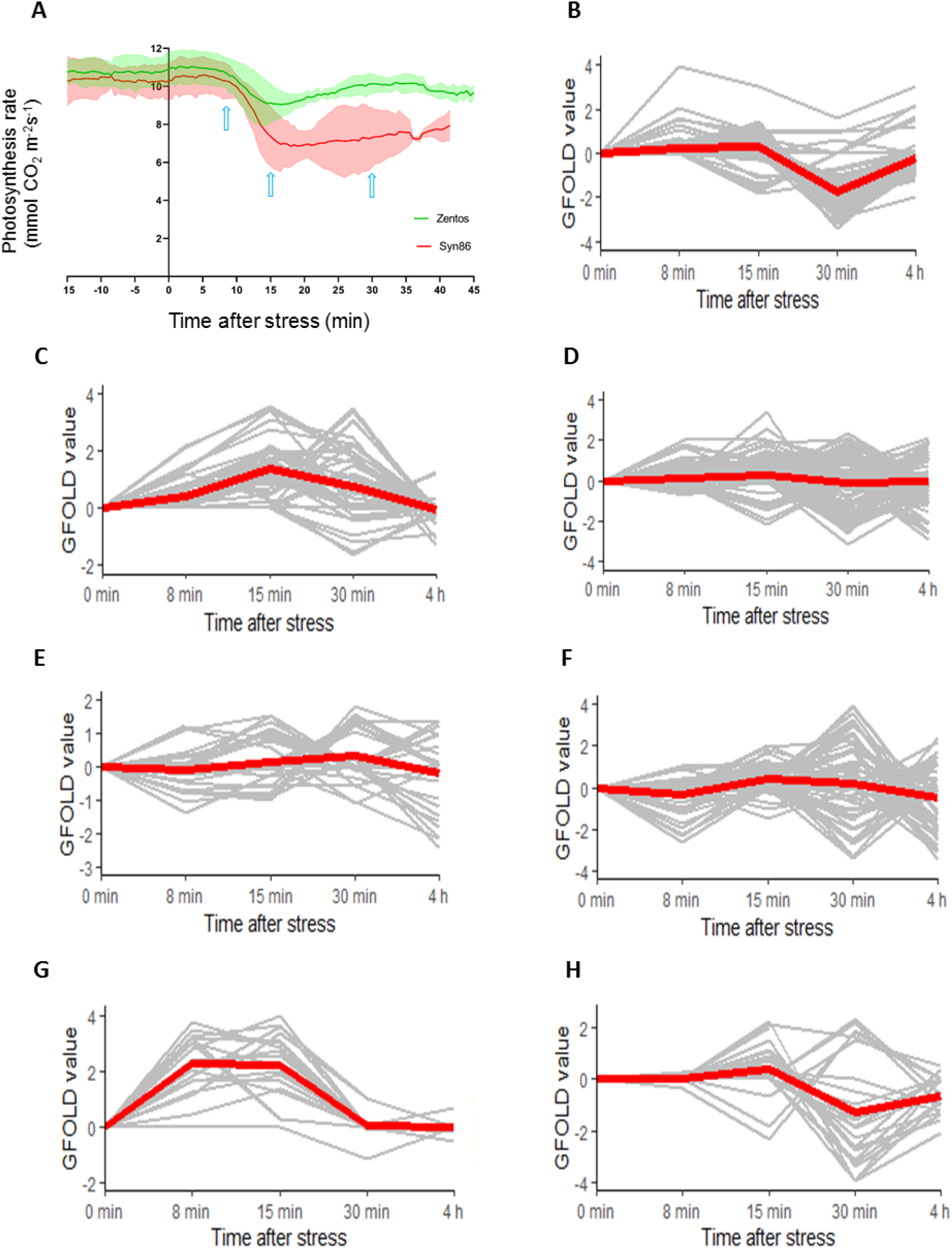
Photosynthesis rate curve (A) and GFOLD values of time course relative expression of some selected gene ontologies from the contrasting genotypes studied during the osmotic stress phase: photosynthesis-related (B), calcium binding (C, D), oxidative stress response (E, F) and xyloglucan:xyloglucosyl transferase activity transcripts (G, H) are shown in Syn86 (B, D, F and H) and Zentos (C, E and G). In each expression profile frame, the gray lines show the time course expression pattern of each gene and the red line is a LOESS (locally estimated scatterplot smoothing) curve that represents the expression tendency of the cluster of genes. The photosynthesis rate curve was adapted from Dadshani (2018), where the shadows represent the standard deviation of the measurements and the time points selected for the transcriptomic analysis are highlighted with blue arrows.

Homogeneous seedlings adapted to hydroponics conditions were used for sampling leaves under control conditions and a salt stress treatment of 100 mol m^−3^ NaCl. The detailed procedures of the hydroponic systems are described in Dadshani (2018) and Oyiga *et al*. (2016) for osmotic and ionic phase experiments, respectively. Osmotic stress conditions were sampled in the photosynthesis turning time points identified at 8, 15 and 30 min (Fig. 1A) and at 4 h ASE whereas control conditions were sampled at 0, 30 min and 4 h in plants grown in hydroponic boxes without NaCl. Samples for the ionic stress conditions were collected at 11 days and 24 days in both salt stressed and unstressed plants. A biologically averaged experiment was conducted where each sampled condition consisted on a pool of five leaves from four plants that were harvested and immediately stored in liquid nitrogen for posterior homogenization. This sampling strategy allowed an exploratory analysis with less amount of sequencing data assuming that most expression measurements from RNA pools are similar to the averages from the individuals that are included in the corresponding pools (Kendziorski *et al*., 2005; Biswas *et al*., 2013).

### MACE reads processing and mapping to the reference genome

The total RNA isolation and the MACE library construction were performed at GenXPro GmbH (Frankfurt, Germany). An Illumina NextSeq 500 system was used to sequence fourteen and eight libraries from the osmotic and ionic stress experiments, respectively. The adapters from the reads were trimmed with Cutadapt (Martin, 2011). The quality control of the prepared libraries was carried out using FastQC (Andrews, 2010) and the short reads with less than 35 bp were removed with Trimmomatic (Bolger *et al*., 2014). The retained reads were aligned to the reference wheat genome assembly version “RefSeq v1.0” (Alaux *et al*., 2018) using Tophat (Trapnell *et al*., 2009). Assemblies of novel transcripts were produced with the prediction tool of Cufflinks (Roberts *et al*., 2011; Trapnell *et al*., 2012). The markdup tool from SAMtools was employed to produce deduplicated alignment files and estimate the amount of read duplication (Li *et al*., 2009).

A new annotation file was elaborated to count reads beyond the predicted 3’-ends of high confidence (HC) and low confidence (LC) gene models (IWGSC, 2018) with the purpose to contribute to gene model improvement and to better estimate gene expression levels. For that, the annotated gene models were extended by 40 % downstream of the predicted 3’-end in the case of intergenic regions greater than 1000 bp but smaller than three times the gene size. When the intergenic distance was larger, the elongated target sequence corresponded to the size of the gene. Then, the stranded option from the featureCounts tool (Liao *et al*., 2014) of the Subread software (Liao *et al*., 2013) was used to count the unique mapped reads assigned to the elongated HC and LC gene models and to novel predicted transcripts. The read count data was normalized to counts per million. An average normalized value of 2.5 across libraries from the same genotype was selected as threshold to define a transcriptomic background aiming to select the genes adequately represented and to reduce the number of low expressed transcripts that might cause sampling noise (Sha *et al*., 2015; Lin *et al*., 2016)

A merged alignment file with all reads from the libraries was generated to compare the number of reads counted using the extended gene models with those counted employing the reference annotation. To exemplify the improvement in transcript quantification with the extended gene models, two windows of 5 Mbp (23 to 28 Mbp coordinates of the chromosomes 5B and 7D) were inspected in the alignment files from two libraries. The online web tool GenomeView (Abeel *et al*., 2012) allowed the visualization of the alignments in the reference genome to observe the coordinates of the reads scored beyond the 3’-end gene boundaries.

### Identification of salt-responsive genes and gene ontology (GO) enrichment analysis

After filtering the low expressed transcripts, salt-responsive genes were identified using the raw count data of fragments as input in the GFOLD (generalized fold change) software (Feng *et al*., 2012). The GFOLD value is a reliable estimator of the relative difference of gene expression which allows the generation of gene rankings that are particularly useful when biological averaging is available (Feng *et al*., 2012). Density plots with log_10_ normalized expression values were generated to assess whether count normalization was appropriate to allow the comparison of the libraries from the same genotype (Klaus and Huber, 2016; available at https://www.huber.embl.de/users/klaus/Teaching/DESeq2Predoc2014.html). The 0 min condition was used as control for both 8 and 15 min ASE, with the assumption that no physiological changes occur in this short period of time under normal conditions. A high absolute GFOLD value indicated greater up- or down-regulation of the genes. Genes with GFOLD values >1 or□<□-1 were considered for further analyses. Venn diagrams were used to summarize the comparisons of the salt-responsive genes across genotypes and time points.

The GO enrichment tool from the STEM software was implemented to distinguish the categories of genes over-represented in the contrasting genotypes (Ernst *et al*., 2005; Ernst and Bar-Joseph, 2006) using the available GO category assignation in the annotation version RefSeq v1.0 (Alaux *et al*., 2018). Only the gene categories from the transcriptomic background of each genotype were retained in the analysis (Timmons *et al*., 2015). A Bonferroni multiple hypothesis correction test was implemented, thus GO terms with a corrected p-value < 0.001 and < 0.005 were considered as over-represented during the osmotic and the ionic phases, respectively. Over-represented categories during the osmotic phase were selected to graphic the time course expression profiles of the corresponding genes. A LOESS (locally estimated scatterplot smoothing) curve was used to represent the expression tendency of the clusters of genes.

### Identification of potential QTGs

The identification of QTGs was done by localizing salt-responsive genes revealed with the transcriptomic analysis within the LD blocks of the corresponding QTL. Thus, adjacent markers in strong LD (R^2^ ≥ 0.8) with the significant single nucleotide polymorphism (SNP) were assigned to one LD block (Cirilli *et al*., 2018). Polymorphic markers in the contrasting genotypes Altay2000 and Bobur were chosen from the association mapping analysis. Afterwards, the positions of the LD blocks in the reference genome sequence RefSeq v1.0 were determined according to the IWGSC RefSeq v1.0 BLAST results of the SNPs-flanking sequences (Alaux *et al*., 2018). Consequently, the LD block coordinates enabled the identification of genes whose transcripts showed up- or down-regulation upon salt treatment and were recognized as potential candidate genes operating within the QTL. The wheat RNA-seq atlas expVIP was used to compare the expression of the genes from the LD blocks with the expression determined in other abiotic stress experiments in the species (Borrill *et al*., 2016).

## Results

### Sequence processing and reference genome mapping

To identify up- or down-regulated transcripts during the salt stress response the samples collected were sequenced using the MACE approach. An overview of the sequence processing and reference genome mapping across both osmotic and ionic stress experiments is presented in the Table 1. The sequencing process yielded a higher number of total and duplicated reads in the eight libraries from Altay2000 and Bobur than in the 14 libraries from the osmotic stress genotypes. The exclusion of less amount of reads after the quality control filtering and a greater average mapping efficiency were observed in the ionic stress libraries compared to those from the osmotic stress experiment (Table 1; see details in Supplementary Table S1).

**Table 1.** Summary of MACE libraries processing and reference genome mapping (mean ± standard deviation).

|  | **Osmotic phase** | **Ionic phase** |
| --- | --- | --- |
| **Libraries** | 14 | 8 |
| **Total millions of reads** | 5.0 ± 0.7 | 9.0 ± 2.4 |
| **Reads excluded after QC<sup>a</sup> (%)** | 7.8 ± 1.6 | 2.6 ± 0.3 |
| **Mapping efficiency (%)** | 83.6 ± 1.2 | 93.0 ± 0.3 |
| **Millions of mapped reads** | 3.8 ± 0.5 | 8.2 ± 2.2 |
| **Multiple aligned reads (%)** | 21.7 ± 3.3 | 19.7 ± 1.7 |
| **Millions of unique mapped reads** | 3.0 ± 0.4 | 6.6 ± 1.8 |
| **Reads after deduplication (%)** | 62.2 ± 3.7 | 53.9 ± 3.2 |

<sup>a</sup>QC: Quality control

The use of the reference annotation file scored 83% of the total number of unique mapped reads while with the extended gene models 88% of the reads were counted. Therefore, with the extended annotation an additional amount of ca. 5 million reads were detected in 12019 genes (see Supplementary Table S2) which accounted for 4.5 % of the gene models predicted in the RefSeq v1.1 genome annotation (IWGSC, 2018). The observation of 10 Mbp from two alignment files allowed the identification of five genes extended from 186 to 470 bp (Table 2). Two of these genes also show a prolonged 3’-end according to RNA-seq data from Pingault *et al*. (2015) visualized in the RefSeq v1.0 genome browser (Alaux *et al*., 2018) (Table 2).

**Table 2.** Comparison of the reads scored with the reference annotation (R) and with extended gene models (E).

| **Gene<sup>a</sup>** | **Reads scored (R/E)** |  | **Extension**<br>**size (bp)** |
| --- | --- | --- | --- |
|  | **Library 1<sup>b</sup>** | **Library 2** |  |
| TraesCS5B02G027100 | 1/9 | 2/3 | 242 |
| TraesCS5B02G028200 | 27/28 | 23/27 | 196 |
| **TraesCS7D02G049200** | 1/2 | 5/16 | 186 |
| TraesCS7D02G062000LC | 5/17 | 4/9 | 368 |
| **TraesCS7D02G051200** | 4/107 | 8/268 | 470 |

<sup>a</sup> Genes in bold letters also show a prolonged 3'-end according to RNA-seq data from Pingault et al. (2015)
<sup>b</sup> Library 1 is from Altay2000 at 11 days under control and Library 2 is from Altay2000 at 24 days of stress without deduplication.

### Identification of salt-responsive genes

To compare the level of expression of genes in response to salt stress in the osmotic and ionic phases, the GFOLD tool was used to identify salt-responsive genes in the two tolerant and two susceptible genotypes studied. The overlapping densities from Syn86 and Zentos indicated an appropriate homogeneity of the sequencing depth from the libraries and an adequate expression normalization (Supplementary Fig. S1). Differently, a greater mean of the expression values was observed at 24 days ASE when compared to the mean in the other time points (Supplementary Fig. S1). This type of distribution of the expression is an indicator of high levels of PCR duplication of reads and therefore the deduplicated alignment files were used for the differential expression analysis at the ionic phase. The density plots produced after deduplication revealed a better homogeneity among samples from the same genotype (Supplementary Fig. S1). The removal of low expressed transcripts leaded to the reduction of the number reads for the differential expression analysis by 3.1% ± 0 3.1% ± 0.4, 3.2% ± 0.2, 3.2% ± 0.6 and 3.9% ± 1.1 for the genotypes Syn86, Zentos, Bobur and Altay2000, respectively.

The differential expression analysis showed a reduced variability among genotypes (mean ± standard deviation) concerning the percentage of identified novel transcripts (4.5 ± 0.4%) (see genome coordinates in Supplementary Table S3), LC (4.6 ± 0.6%) and HC (90.9 ± 1.0%) gene models. The D subgenome contained the greatest percentage of salt-responsive genes (35.8 ± 1.7%) followed by subgenomes B (31.4 ± 1.1%) and A (31.3 ± 1.9%) and unplaced superscafolds (1.5 ± 0.4%) (see all salt-responsive genes in Supplementary Table S4).

### Comparative analysis of the osmotic stress response

To better understand the early plant reaction to salt exposure, the comparative transcription profiling at osmotic stress phase was performed. This analysis revealed up- and down-regulated salt-responsive genes across the four time points studied. The greatest amount of them was observed at 15 min ASE in Zentos and at 30 min in Syn86, whereas the lowest number of salt-responsive genes was identified at 8 min ASE in both genotypes (Fig. 2B,C). Thirty eight and fourteen genes were differentially expressed simultaneously across all the time points in Syn86 and Zentos, respectively (Fig. 2B,C). The distribution of up and down-regulated genes across the time points of the osmotic stress experiment are presented in the Fig. 3A. Zentos had the highest number of up- and down-regulated genes at 15 min and 4 h ASE, respectively. In contrast, Syn86 had the highest number of up-regulated genes at 4 h and of down-regulated at 30 min. In total, Zentos showed 75% of up-regulated genes while Syn86 had 60%.

**Fig. 2.**
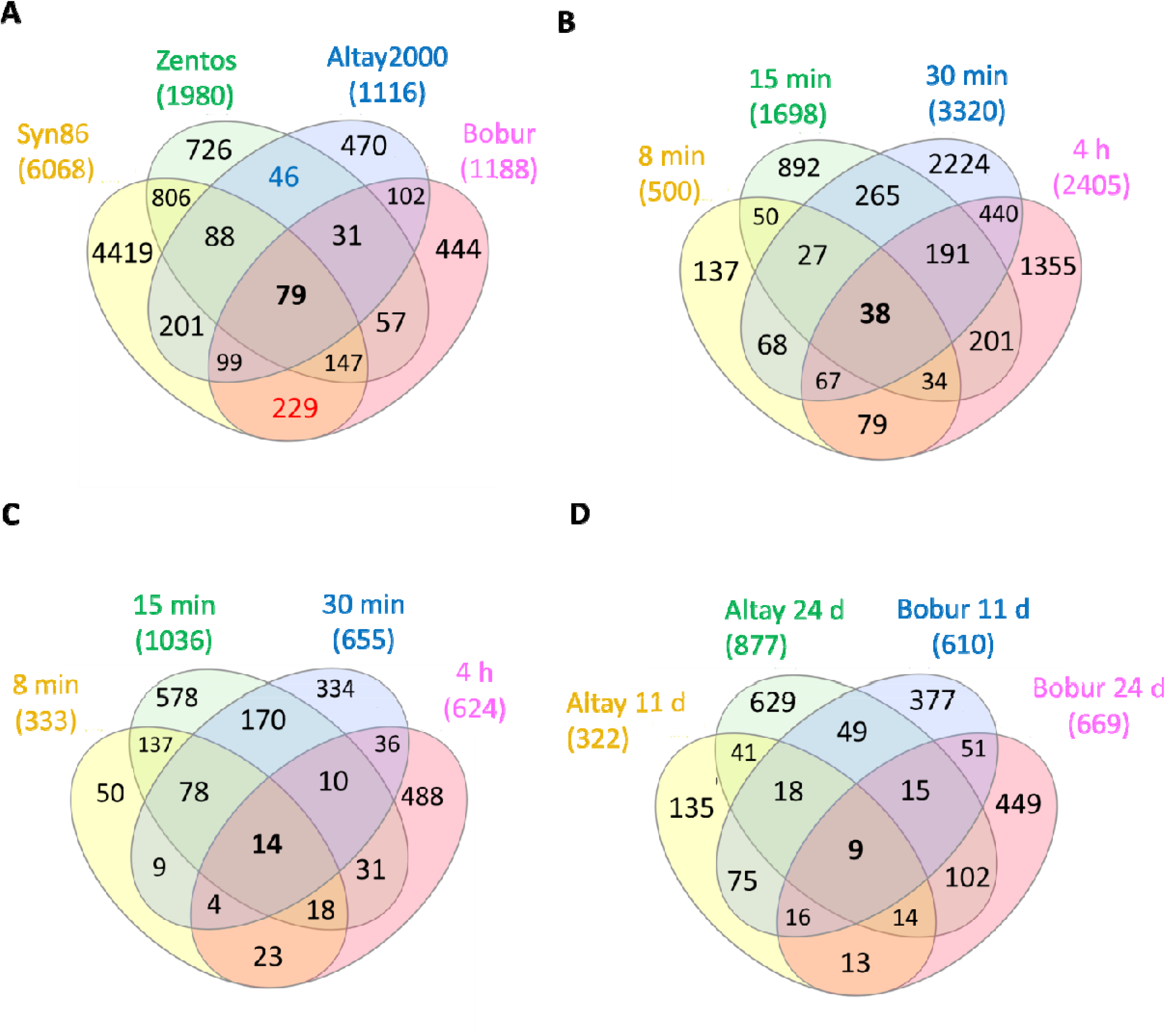
Venn diagrams of the salt-responsive genes in the contrasting genotypes studied. The total amount of genes in each genotype and/or time point are shown above each diagram. (A) Diagram with the four genotypes. The blue number represents the genes shared by the tolerant genotypes while the red number indicates the genes shared by the salt-susceptible; (B) diagram of the salt-responsive genes in Syn86 by time point; (C) diagram of the salt-responsive genes in Zentos by time point; (D) diagram of the salt-responsive genes in the two sampled days from the ionic stress phase.

**Fig. 3.**
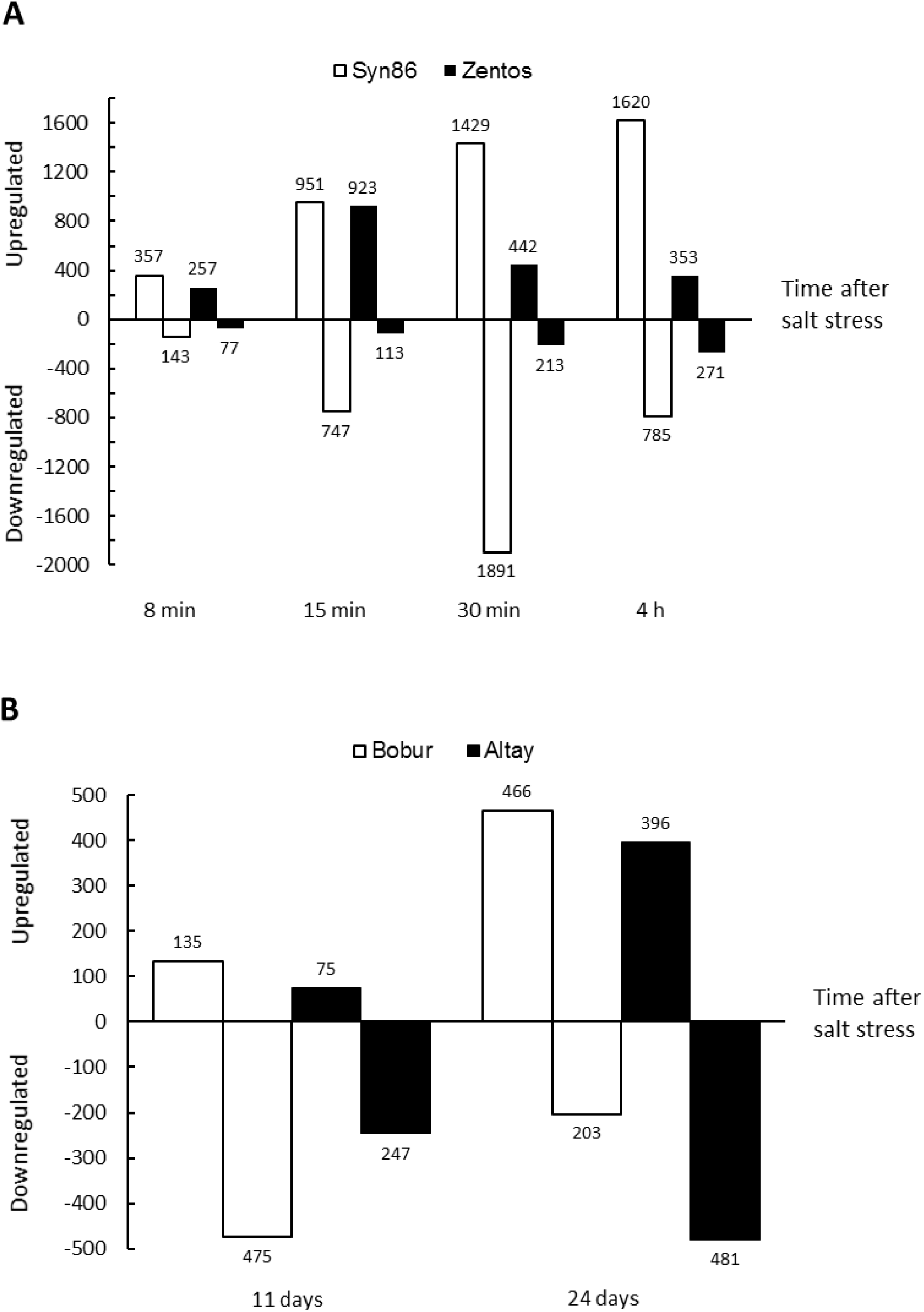
Distribution of up- and down-regulated salt-responsive genes across stress time points. (A) Osmotic phase and (B) ionic phase time points.

The GO enrichment analysis allowed to compare the over-represented gene categories among the up- and down-regulated genes identified in each genotype and time point. Highlighted in the heatmaps are the 24 and 18 ontology terms that were exclusively up- and down-regulated, respectively (Fig. 4A,B). Among the up-regulated categories, the over-representation of response to wounding genes and tryptophan synthase activity were recognized in the susceptible genotype, whereas in Zentos the calcium binding category was identified. Defense response to fungus and bacterium, transcription factor activity and protein kinase coding genes were over-represented and up-regulated in both genotypes (Fig. 4A). The down-regulated categories revealed the over-representation of spermine and spermidine biosynthesis and antioxidant activity genes in Syn86, while the lipid binding category was observed in both genotypes (Fig. 4B).

**Fig. 4.**
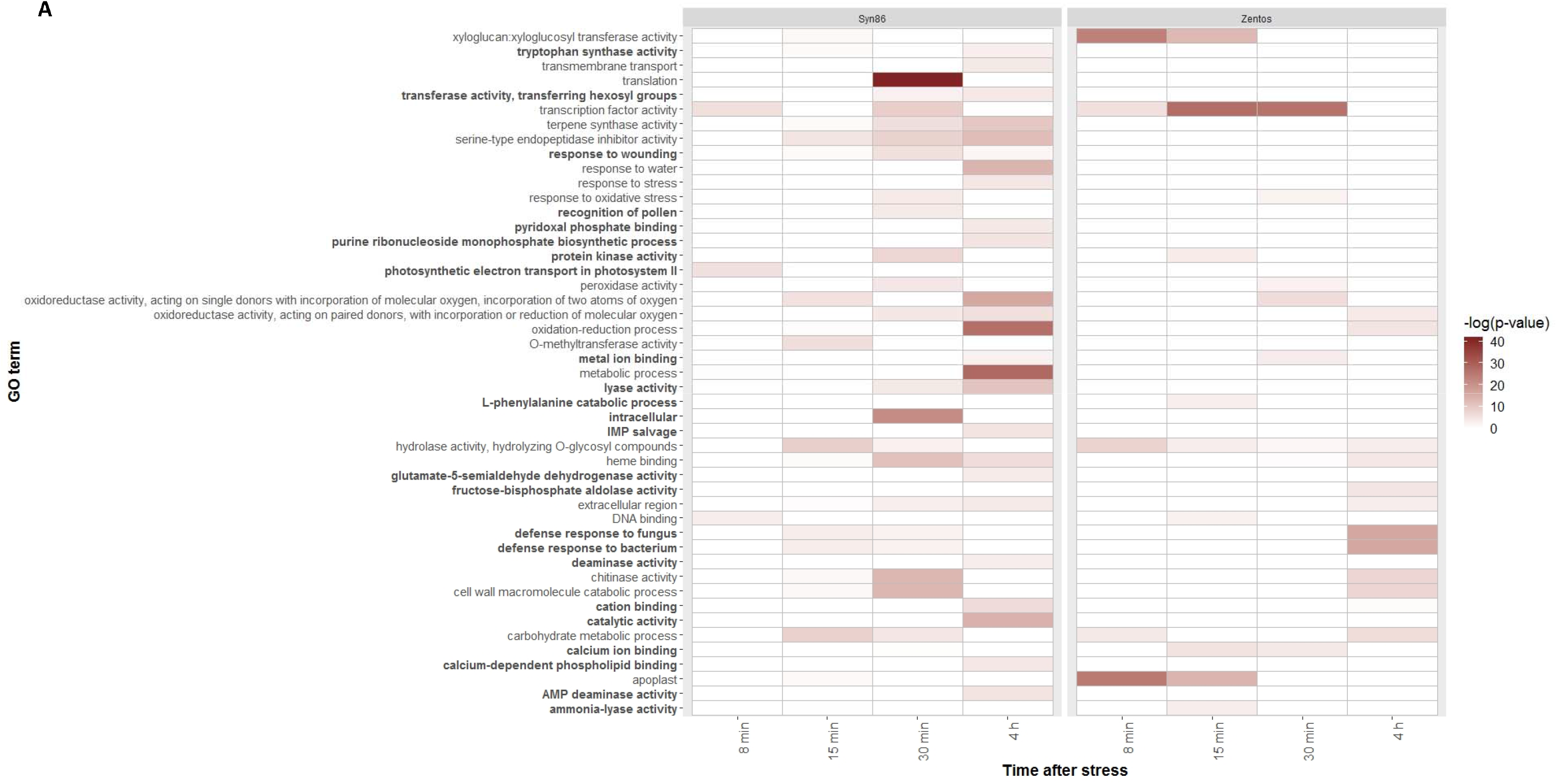
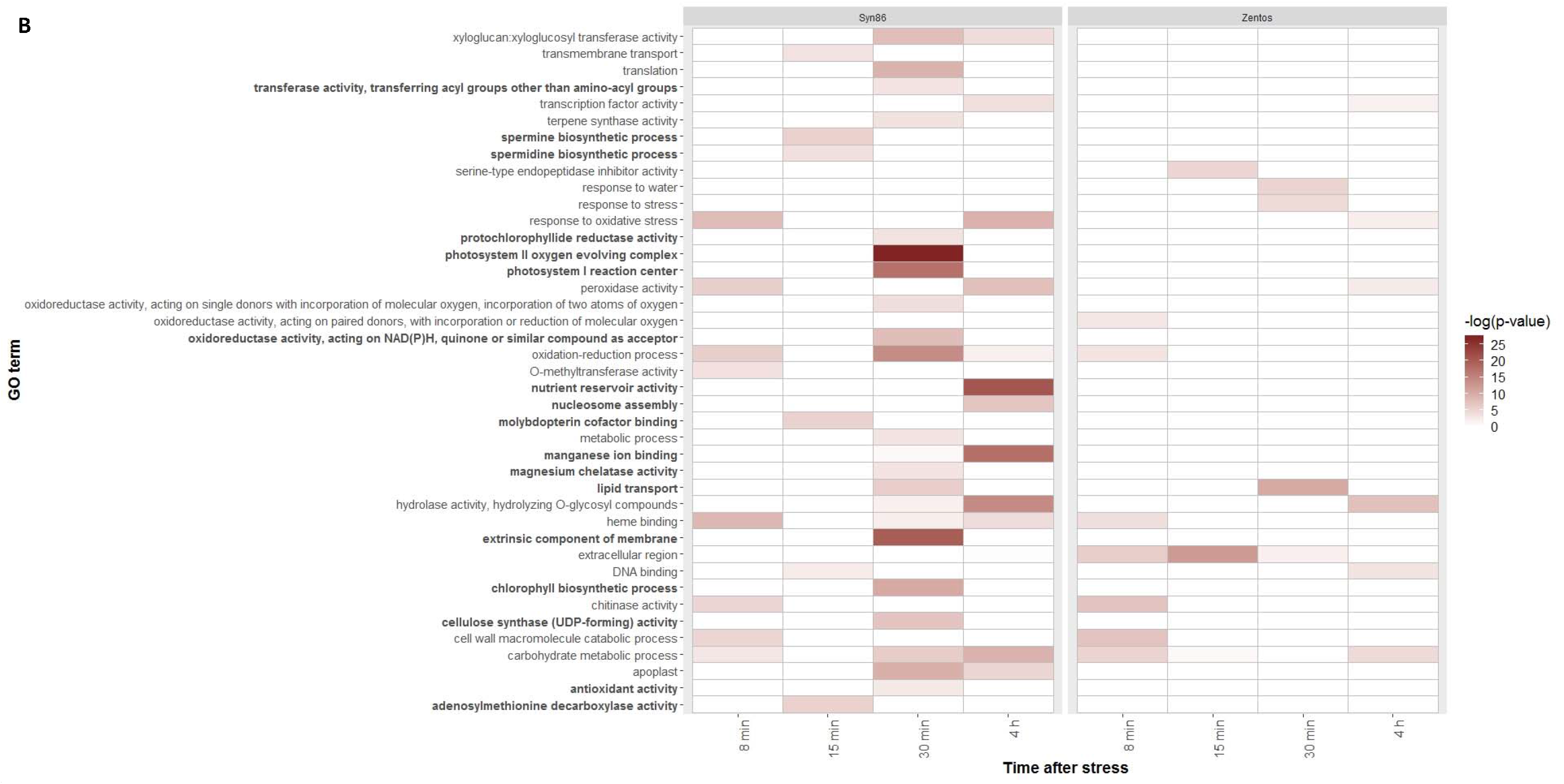
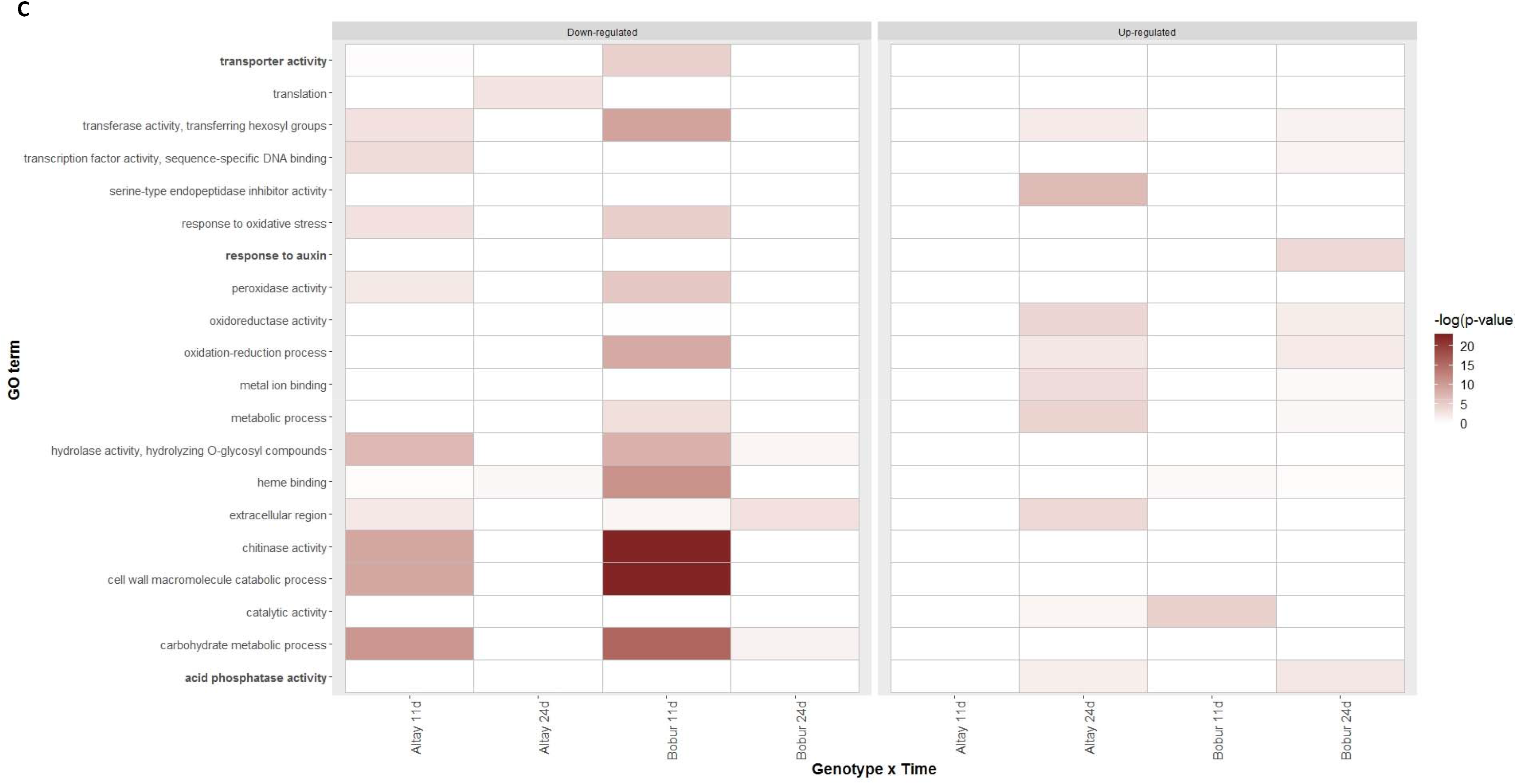
GO terms over-represented during the salt stress response. (A) Up-regulated and (B) down-regulated categories identified in the four stress time points sampled during the osmotic phase; (C) up- and down-regulated categories observed in the two stress time points from the ionic phase. Bold ontologies are categories specific for each heatmap. The –log_10_ transformation of the corrected p-values highlights the categories with greater significance that are therefore better over-represented.

Some gene ontologies such as xyloglucan:xyloglucosyl transferase activity and oxidative stress were over-represented in both genotypes and revealed differential expression profiles. The calcium binding category also showed differential expression profiles in the genotypes, but unlike Zentos, this term was not over-represented in Syn86 (p-value > 0.05) even though more genes showed differential expression (129 vs 50 genes). Additionally, ontology terms related to photosynthesis were over-represented in both up- and down-regulated genes from Syn86 (Fig. 4A,B). Because of the relevance of these categories in the osmotic stress response and the link of oxidative stress, calcium binding and photosynthesis categories with the photosynthetic response under salt stress, a comparison of the expression profiles in the contrasting genotypes will be presented in the next section.

### Time course of gene expression and photosynthesis rate during the osmotic phase

A comparison of the expression profiles of the salt-responsive calcium binding, oxidative stress response and xyloglucan:xyloglucosyl transferase activity genes was performed to understand the time course of gene expression differences of these categories in the contrasting genotypes (Fig. 1). The expression of 101 photosynthesis-related genes is only shown in the susceptible genotype (Fig. 1B) since the related categories were not over-represented in the tolerant genotype. The up-regulation of eight electron transport in PSII (photosystem II) genes was observed at 8 min ASE when the photosynthesis rate starts to decrease in Syn86 (Fig. 1A). When the photosynthesis rate showed recovery but was still inhibited (Fig. 1A), 91 transcripts from both photosystems I and II were down-regulated at 30 min with relative expression values ranging from −1.1 to −3.4 (Fig. 1B).

The LOESS curve from the 50 salt-responsive calcium binding genes of the tolerant genotype revealed a gene up-regulation tendency at 15 min. Thirty-four transcripts were identified in this time point with relative expression values ranging from 1.0 to 3.4 (Fig. 1C). From these genes, 32 contained an EF hand calcium binding domain. The susceptible genotype presented heterogeneous expression patterns for 129 salt-responsive genes of this category. In this genotype, the greatest amount of calcium binding genes (40) were down-regulated at 30 min with GFOLD values ranging from −1.0 to −3.1 (Fig. 1D). The majority from these genes (26) were components of the oxygen-evolving complex from the PSII (Wang *et al*., 2019). This result was also in line with the suppressed photosynthesis rate of Syn86 at this time point (Fig. 1A). Furthermore, 35 and 21 additional genes from this category were up-regulated at 30 and 15 min, respectively.

On the other hand, 33 salt-responsive genes from the oxidative stress response category were identified in Zentos. Eight and 10 of them were up-regulated and showed relative expression values lower than 2.5 at 15 and 30 min ASE, respectively. The down-regulation of eight genes was observed at 4 h with expression values ranging from −1.0 to −2.4 (Fig. 1E). In contrast, Syn86 contained 59 genes displaying heterogeneous expression patterns with greater relative expression values than Zentos (Fig. 1F). Thus, 11, 17 and 22 genes were down-regulated at 8, 30 min and 4 h, respectively, while 21 transcripts at 30 min and 11 genes at 4 h were up-regulated. The expression values from the down-regulated transcripts fluctuated from −1.0 to − 3.5 and the up-regulated genes revealed a GFOLD value range from 1.0 to 4.2. The greatest number of salt-responsive oxidative stress genes observed at 30 min (38 transcripts), which included both up- and down-regulated transcripts, agrees as well with the inhibited photosynthetic activity observed in Syn86 in this time point (Fig. 1A).

Finally, all the salt-responsive cell wall genes corresponded to the xyloglucan:xyloglucosyl transferase activity category. Eighteen genes were identified in Zentos from which 14 showed up-regulation both at 8 and 15 min with GFOLD values ranging from 1.1 to 4.0 (Fig. 1G). Otherwise, 24 genes from Syn86 were observed in this category where the LOESS curve highlighted the down-regulation of 16 transcripts at 30 min (Fig. 1H). The relative expression values of the down-regulated transcripts ranged from −1.0 to −4.3.

### Comparative analysis of the ionic stress response

To better understand the later phase of plant reaction to salt exposure, the comparative transcription profiling at ionic stress phase was performed with the expression values calculated after deduplication. This analysis revealed the fewest amount of salt-responsive genes in Altay at 11 days (Fig. 2D). The simultaneous differential expression of nine genes was identified across genotypes and time points (Fig. 2D). The Fig. 3B shows the distribution of up and down-regulated genes. At 24 days ASE Bobur showed a greater proportion of up-regulated than of down-regulated ones, whereas the opposite pattern with greater amount of down-regulated transcripts was found in Bobur at 11 days ASE and in Altay2000 at both time points (Fig. 3B). In total, Altay2000 and Bobur contained 61% and 53% of down-regulated genes, respectively.

The Fig. 4C summarizes the GO enrichment analysis at the ionic stress phase which is separated by the up-and down-regulated genes in the two genotypes and the two time points. Three GO terms specific for this stress phase were identified and half of the enriched categories shared the same stress effect in the two genotypes (Fig. 4C). For instance, transferase activity, chitinase activity and response to oxidative stress were down-regulated in both genotypes (Fig. 4C). On the other hand, translation and transcription factor activity terms were down-regulated in the tolerant genotype while metal ion binding was up-regulated. The up-regulation of the response to auxin category and the down-regulation of transporter activity were observed in the susceptible genotype (Fig. 4C).

### Comparative analysis of osmotic and ionic stress responses

The implementation of a transcriptomic approach allowed the comparison of the salt stress response during the osmotic and ionic phases in the analyzed genotypes. Syn86 was the genotype presenting the highest amount of salt-responsive genes, from three to five times more genes than the three cultivars. From all the differentially expressed genes, 79 were stress responsive in the four genotypes while 46 and 229 transcripts were expressed in both tolerant and both sensitive genotypes, respectively (Fig. 2A).

A total of 17 GO terms were over-represented in both the osmotic and ionic phases of salt stress (Fig. 4). The translation category appeared down-regulated in the salt-sensitive Syn86 at the osmotic phase and the tolerant Altay2000 at the ionic phase. The serine-type endopeptidase inhibitor activity term presented opposite relative expression values in the tolerant genotypes of both salt stress phases. These genes were down-regulated in the tolerant genotype and up-regulated in the salt-sensitive one during the osmotic phase. On the contrary, this category showed up-regulation in the salt-tolerant genotype at the ionic stress phase. The response to oxidative stress category was both up- and down-regulated in the contrasting genotypes from at osmotic stress phase while it was only down-regulated in both genotypes studied during the ionic phase.

### Identification of candidate QTGs

To unravel candidate QTGs that might contain alleles controlling salt stress-related traits, salt-responsive transcripts within the LD blocks harboring markers with significant phenotypic effect were identified and analyzed. The Table 3 presents the candidate QTGs from two QTL identified in an association panel (Oyiga *et al*., 2018) and an AB-QTL mapping population (Dadshani, 2018) on the chromosome 2A. A 36 Mbp LD block was covered by the QTL interval detected by the marker RAC875_c38018_278. This marker showed an effect on shoot fresh weight after salt stress in the association mapping panel. Two differentially expressed genes were found in this region, one salt-responsive in the sensitive genotype and one in the tolerant (Table 3). Among them, TraesCS2A02G395000 showed the strongest stress response since the gene coding an oxoglutarate/iron-dependent dioxygenase was suppressed in the salt-susceptible genotype with an expression value of −2.4. On the AB-QTL mapping population, a 9 Mbp LD block constituted the QTL interval from the marker BS00041707_51. A marker-trait association with kernel weight under stress was discovered with this SNP. This region included two up-regulated genes in Syn86 with similar relative expression values: TraesCS2A02G327600 which codes for a copper amine oxidase and TraesCS2A02G331100 which codes for an amino acid transporter. The stress effect on the expression was confirmed in other abiotic stress studies for the four candidate QTGs determined in both intervals (Table 3).

**Table 3.** Differentially expressed genes in LD blocks of markers with effect on salt stress-related traits in the Chr 2A.

| Marker<sup>a</sup> | LD interval and size<br>(cM / Mbp) | No. SNPs in interval<br>(No. in LD) | Gene relative expression<sup>d</sup> | Annotation<sup>e</sup> | Abiotic stress<br>effect<sup>f</sup> |
| --- | --- | --- | --- | --- | --- |
| RAC875_c38018_278<sup>b</sup> | 109.51-112.14= 2.63/<br>612-648=36 | 7 (5) | TraesCS2A02G389400: **1.5**<br>TraesCS2A02G395000: -2.4 | Leucine zipper, homeobox-associated<br>Oxoglutarate/iron-dependent dioxygenase | 1) ↑, 2) ↑<br>1) ↓ |
| BS00041707_51<sup>c</sup> | 105.53-108.46=2.93/<br>556-565=9 | 34(5) | TraesCS2A02G327600: 2.1<br>TraesCS2A02G331100: 1.8 | Copper amine oxidase<br>Amino acid transporter, transmembrane domain | 1) ↑, 2) ↑<br>1) ↑ |

<sup>a</sup> Marker names according to Wang et al. (2014)
<sup>b</sup> From mapping study (Oyiga *et al.*, 2018)
<sup>c</sup> From mapping study (Dadshani, 2018)
<sup>d</sup> GFOLD values from tolerant genotypes are bold and from the susceptible are in italics.
<sup>e</sup> Based on the Interpro results from the RefSeqv1.0 annotation (Alaux *et al.*, 2018).
<sup>f</sup> Abiotic stress response based on studies deposited in the wheat expression atlas expVIP (Borrill *et al.*, 2016). 1) Drought and heat (Liu *et al.*, 2015) and 2) cold (Li *et al.*, 2015). The direction of the arrows indicates the stress effect on expression, ↑ when the gene is up-regulated and ↓ when is down-regulated.

## Discussion

### Novel regions with transcription in wheat identified with MACE sequencing

The bioinformatic pipeline implemented detected salt-responsive genes in the contrasting wheat genotypes studied. This pipeline has also been applied in recent transcriptomics studies with RNA pools from plants and the real-time PCR validations analyses have confirmed its high accuracy (Qiao *et al*., 2017; Vidya *et al*., 2018). Besides uncovering the dynamic transcriptomic response during salt stress, the MACE-derived sequence analysis conferred evidence of two type of novel regions with transcription in wheat. Firstly, the differential expression analysis assigned a putative role in the salt stress response to *in silico* predicted novel transcripts. These novel salt-responsive transcripts might enrich the wheat variable pangenome that represents 39% of the pangenome according to the analysis of the whole genome of 18 cultivars (Montenegro *et al*., 2017). Secondly, the detection of unpredicted 3’-ends of genes gave indication of longer gene transcription contributing to the improvement of the current gene models (IWGSC, 2018). These 3’-end-extended transcripts might reveal that some current gene model predictions were based on transcripts with incomplete read coverage in the region possibly due to low expression levels in previous transcriptomic experiments (Roberts *et al*., 2011). The genes identified with extended transcription can be included in computational prediction approaches to better define gene structures (Inatsuki *et al*., 2016; Tzfadia *et al*., 2018). The real-time PCR validation of both novel transcripts and 3’-ends is necessary to confirm the transcription of these regions.

### Comparative transcriptomic and time course of photosynthesis rate during the osmotic phase of stress

The adequate expression normalization of the libraries from Syn86 and Zentos allowed to compare the gene expression values of the same genotype at the different conditions (Klaus and Huber, 2016; available at https://www.huber.embl.de/users/klaus/Teaching/DESeq2Predoc2014.html). The osmotic stress experiment revealed the early up-regulation and the posterior down-regulation of photosynthesis-related transcripts in the susceptible genotype. The up-regulation at 8 min of the electron transport in PSII category can be linked to the over-excitation of this system which leads to an increase in the generation of ROS (Parihar *et al*., 2015; Foyer, 2018). The down-regulation of photosynthesis-related genes at 30 min ASE might be a consequence of excessive ROS accumulation that inhibits the repair of photodamaged PSII at both transcriptional and translational levels (Allakhverdiev *et al*., 2002; Murata *et al*., 2007; Saibo *et al*., 2009; Queval and Foyer, 2012). Nevertheless, the transcriptional suppression of photosynthesis is not observed at 4 h of stress (Fig. 1A), which indicates that there are mechanisms that allow the plant to recover the normal expression level of photosynthesis-related genes.

The reduced oxidative stress response of Zentos can be attributed to a restrained ROS production in the tolerant genotype which might have a stimulating effect on the growth under stressful conditions (Queval and Foyer, 2012). The reduced photosynthesis inhibition of this genotype can be therefore linked to a lower oxidative damage of the photosynthetic apparatus. Differently, the susceptible genotype revealed both the down- and up-regulation of genes implicated in oxidative damage protection with greater relative expression values than the tolerant genotype (Fig. 1E,F). These results indicate that salt stress exerted a stronger effect on the oxidative damage protection system of Syn86 at the transcriptional level which supports its greater photosynthesis inhibition.

The over-representation of genes coding for calcium binding proteins at 15 min in Zentos agrees with an earlier timing of calcium and ROS signaling proposed for salt-tolerant genotypes (Ismail *et al*., 2014). These molecules interact to regulate salt stress response (Ismail *et al*., 2014; Parihar *et al*., 2015; Köster *et al*., 2018). A delayed Ca^2+^/ROS signaling will lead to the activation of jasmonic acid signaling pathway that will culminate in cell death. Differently, an earlier activation of calcium-and ROS-dependent signaling induces the constraint of jasmonic acid signaling through the activation of the abscisic acid signaling pathway (Ismail *et al*., 2014). We can infer that the calcium binding genes that exhibited an early up-regulation in Zentos may play a role in activating the signaling pathway leading to abscisic acid accumulation to stimulate growth under stress conditions (Ergen *et al*., 2009). In contrast, in the susceptible genotype the calcium binding up-regulation was delayed and occurred at 30 min ASE which is consistent with the described model (Ismail *et al*., 2014). In addition to a delayed calcium binding up-regulation, the salt-driven suppression of calcium binding genes related to photosynthesis was observed as well. This result is also linked to the photosynthesis inhibition observed in Syn86.

The present study revealed as well the differential response of the xyloglucan:xyloglucosyl transferase activity term in the contrasting genotypes. The greater transcription observed in the tolerant genotype might enable plant growth under stress and might be beneficial for cell wall strengthening, the prevention of excessive water loss and to maintain turgor pressure because of the biosynthesis of xyloglucan in the cell wall (Eckardt, 2008; Le Gall *et al*., 2015). On the other hand, the down-regulation of these genes in the salt-sensitive genotype might be linked to the inhibition of cell expansion and cell wall synthesis that limit plant growth under stress conditions (Wang *et al*., 2018). The synergy of the described transcriptional events might be crucial for the contrasting salt stress response of Syn86 and Zentos during the osmotic phase.

### Transcriptomic response during the ionic phase

The distribution of the expression values at the ionic phase insinuated a high amount of PCR duplication in the libraries at 24 days ASE which coincided with the higher amount of reads sequenced in this time point in both genotypes. The use of deduplicated read alignments was therefore chosen to reduce false positives in the identification of differentially expressed genes and to allow a better comparability among libraries. PCR duplication leads to the overestimation of transcript abundance because the number of reads is biased in the libraries (Klepikova *et al*., 2017).

Among the over-represented transcripts identified, the ionic stress effect on the expression of metal ion binding genes was in accordance with their role in oxidative stress response and antioxidant activity in the photosynthetic electron transport system because of their redox potentials (Palma *et al*., 2013; Yruela, 2013). Therefore, the up-regulation of this mechanism might counteract the oxidative damage in the tolerant genotype at 24 ASE since at 11 days oxidative stress response activity was down-regulated. The observed down-regulation of transporter activity in Bobur suggests two possibilities. First, it might be a strategy of the susceptible genotype to avoid Na^+^ transport and accumulation in the photosynthetic tissues but also may point out at a potential harmful effect of this mechanism for the uptake of non-toxic elements relevant for plant growth when the expression of non-selective cation channels is affected (Assaha *et al*., 2017). A second possibility might be the down-regulation of transporters responsible for Na^+^ exclusion from leaves that will lead to genotype susceptibility (Cotsaftis *et al*., 2012; Wu, 2018).

The similar stress response of some GO terms observed in the contrasting genotypes at the ionic stress phase supports the finding that some earlier transcriptional responses might present stronger differences and might cause a greater impact in the contrasting acclimation response of the genotypes to long-term salt stress (Julkowska and Testerink, 2015). Nevertheless, it is also possible that when similar pathways are salt-responsive in both genotypes the difference might lie in the magnitude of the expression to affect the differential response.

### Comparative transcriptomic analysis of osmotic and ionic phases of salt stress

The stress effect on mechanisms related with protein synthesis and breakdown was identified in the comparative analysis of both stress phases. An accumulation of aberrant proteins in cells can result from stress-related ROS damage which can lead to the transient suppression of the *de novo* synthesis of proteins and the intracellular protein degradation by proteases (Kidrič *et al*., 2014; Zhu, 2016; Robles and Quesada, 2019). The differential response of the translation and the serine-type endopeptidase inhibitor categories across the stress phases suggests that the regulation of these mechanisms are stress stage specific.

### Identification of candidate QTGs

Among the two salt-responsive genes observed in the QTL interval from the association mapping analysis, the oxoglutarate/iron-dependent dioxygenase showed the strongest down-regulation. This gene superfamily with a wide functional diversification is involved in the biosynthesis of several specialized secondary metabolites responsive to biotic and abiotic stresses (Farrow and Facchini, 2014; Xu and Song, 2017). Therefore, this gene is a strong candidate that can be prioritized for further validation analyses. The AB-QTL mapping interval contained as well two salt-responsive genes including a copper amine oxidase and an amino acid transporter with similar magnitudes of relative expression. The up-regulation of both genes in the sensitive genotype can be linked to the positive phenotypic effect of the allele from Syn86 in the variation of kernel weight under salt stress (Dadshani, 2018). Studies in *Arabidopsis* have shown the involvement of copper amine oxidases in the biosynthesis of nitric oxide which is a signaling molecule that participates in adaptive responses to biotic or abiotic stresses (Neill *et al*., 2002; Wimalasekera *et al*., 2011; Groß *et al*., 2017). On the other hand, there are amino acid transporters up-regulated by salt stress that are involved in the transport of amino acids as proline which accumulates under stress to act as an osmolyte for osmotic adjustment (Hayat *et al*., 2012; Wan *et al*., 2017). The differential expression of the genes in the interval may contribute concomitantly to the phenotypic variation (González-Prendes *et al*., 2017). Only the implementation of a higher resolution mapping approach and functional studies could help to confirm the causality of the selected candidate genes on the trait of interest.

The above mentioned results validate the use of the recent genome assembly to facilitate the integration of genomic and transcriptomic resources to resolve QTL and advance in the understanding of the genetic variation underlying agronomic complex traits in bread wheat (Adamski *et al*., 2018; IWGSC, 2018). This screening approach to target potential functional candidate genes is relevant since the mapping resolution of the studies is limited and the average gene density considering solely HC models is of approximately seven genes per Mbp (IWGSC, 2018). Nevertheless, this strategy can be more robust when expression data of other tissues under salt stress and different time points can be also included.

### Conclusions and future perspectives

To our knowledge this is one of the first studies to combine transcriptomics, bioinformatics, genetics and stress physiology analyses in a systemic approach to obtain a more comprehensive understanding of the salt stress adaptation response in bread wheat at both osmotic and ionic phases. We expect that our results will encourage the wheat research community to perform functional analysis of some prioritized genes. This will lead to a better QTL dissection to finally shed light on novel genes controlling regulatory pathways for salt stress-related traits.

## Supplementary data

**Table S1.** Overview of reads processing and reference genome mapping from all libraries.

**Table S2.** Genes with extended 3’-end and position in the reference genome of the clusters of additional reads scored.

**Table S3.** Reference genome coordinates from the novel salt-responsive transcripts identified in the four genotypes.

**Table S4.** GFOLD values of the salt-responsive genes identified in the time points studied in the four genotypes.

**Fig. S1.** Density plots with the log_10_ normalized expression values of the libraries from the four genotypes studied.

## Data availability

The alignment of reads data generated from the libraries were deposited to the Sequence Read Archive of the National Center for Biotechnology Information with the accession number SUB5849358.

## Acknowledgements

This study was financed by the “Bundesministerium für wirtschaftliche Zusammenarbeit und Entwicklung” via “German Agency for International Cooperation”, Germany (Project 09.7860.1-001.00). DDD was funded by the PhD scholarship call 679 from Colciencias (Science, Technology and Innovation Department from Colombia). The authors would like to acknowledge A. Shresta and S. Parra for the critical reading of the manuscript.

