## Supplementary material for "A systemic approach provides insights into the salt stress adaptation mechanisms of contrasting bread wheat genotypes": Suppl Table S1 and Fig S1

**Table S1a**. Overview of osmotic stress libraries processing and reference genome mapping

| Libraries | | Total reads | Filtered reads after QC[b] (%) | Mapped reads | Mapping efficiency (%) | Unique mapped reads | Multiple aligned reads (%) |
|---|---|---|---|---|---|---|---|
| **Syn86 Control** | **0 min** | 3762527 | 7.1 | 2941718 | 84.2 | 2088265 | 29.0 |
| | **30 min** | 5040446 | 6.6 | 3977669 | 84.5 | 3047297 | 23.4 |
| | **4 h** | 5706140 | 9.2 | 4300722 | 83.0 | 3530248 | 17.9 |
| **Syn86 Stress** | **8 min** | 3854936 | 5.2 | 3025224 | 82.8 | 2257892 | 25.4 |
| | **15 min** | 5049633 | 7.3 | 3862225 | 82.5 | 3080479 | 20.2 |
| | **30 min** | 5262981 | 7.9 | 3998840 | 82.5 | 3281663 | 17.9 |
| | **4 h** | 4255947 | 7.0 | 3291831 | 83.2 | 2632432 | 20.0 |
| **Zentos Control** | **0 min** | 5572114 | 9.4 | 4218244 | 83.6 | 3287693 | 22.1 |
| | **30 min** | 5212469 | 8.5 | 4043075 | 84.7 | 3200059 | 20.9 |
| | **4 h** | 5235429 | 7.8 | 4085748 | 84.7 | 3136558 | 23.2 |
| **Zentos Stress** | **8 min** | 6068132 | 9.7 | 4640926 | 84.7 | 3683923 | 20.6 |
| | **15 min** | 5460661 | 9.7 | 4195570 | 85.1 | 3119621 | 25.6 |
| | **30 min** | 4133679 | 4.6 | 3299731 | 83.7 | 2660847 | 19.4 |
| | **4 h** | 5233986 | 8.8 | 3867463 | 81.0 | 3155938 | 18.4 |
| **Average** | | 4989220.0 | 7.8 | 3839213.3 | 83.6 | 3011636.8 | 21.7 |
| **SD[a]** | | 709964.9 | 1.6 | 506378.5 | 1.2 | 451258.6 | 3.3 |

[a]Standard deviation
[b]Quality control

**Table S1b**. Overview of ionic stress libraries processing and reference genome mapping

| Libraries | | Total reads | Removed after QC (%) | Mapped reads | Mapping efficiency | Unique mapped Reads (UMR) | Multiple aligned reads (%) | UMR counted after deduplication |
|---|---|---|---|---|---|---|---|---|
| **Altay2000** | **11 days** | 8114165 | 2.47 | 7388399 | 93.4 | 5685325 | 23.1 | 2209963 |
| **Control** | **24 days** | 12148620 | 2.51 | 11052250 | 93.3 | 8934228 | 19.2 | 3049629 |
| **Altay2000** | **11 days** | 5123752 | 2.78 | 4633841 | 93.0 | 3654709 | 21.1 | 1544343 |
| **Stress** | **24 days** | 8177854 | 2.73 | 7364984 | 92.6 | 6005757 | 18.5 | 2677389 |
| **Bobur** | **11 days** | 9681088 | 2.97 | 8717906 | 92.8 | 7126152 | 18.3 | 2566285 |
| **Control** | **24 days** | 9913293 | 2.16 | 9079634 | 93.6 | 7376510 | 18.8 | 3001207 |
| **Bobur** | **11 days** | 7204855 | 2.48 | 6537136 | 93.0 | 5241318 | 19.8 | 2149826 |
| **Stress** | **24 days** | 11960398 | 2.42 | 10846554 | 92.9 | 8849873 | 18.4 | 3376956 |
| **Average** | | 9040503.1 | 2.6 | 8202588.0 | 93.1 | 6609234.0 | 19.7 | 2571949.8 |
| **SD** | | 2380609.1 | 0.3 | 2171900.5 | 0.3 | 1816597.3 | 1.7 | 590014.8 |

[a]Standard deviation
[b]Quality control

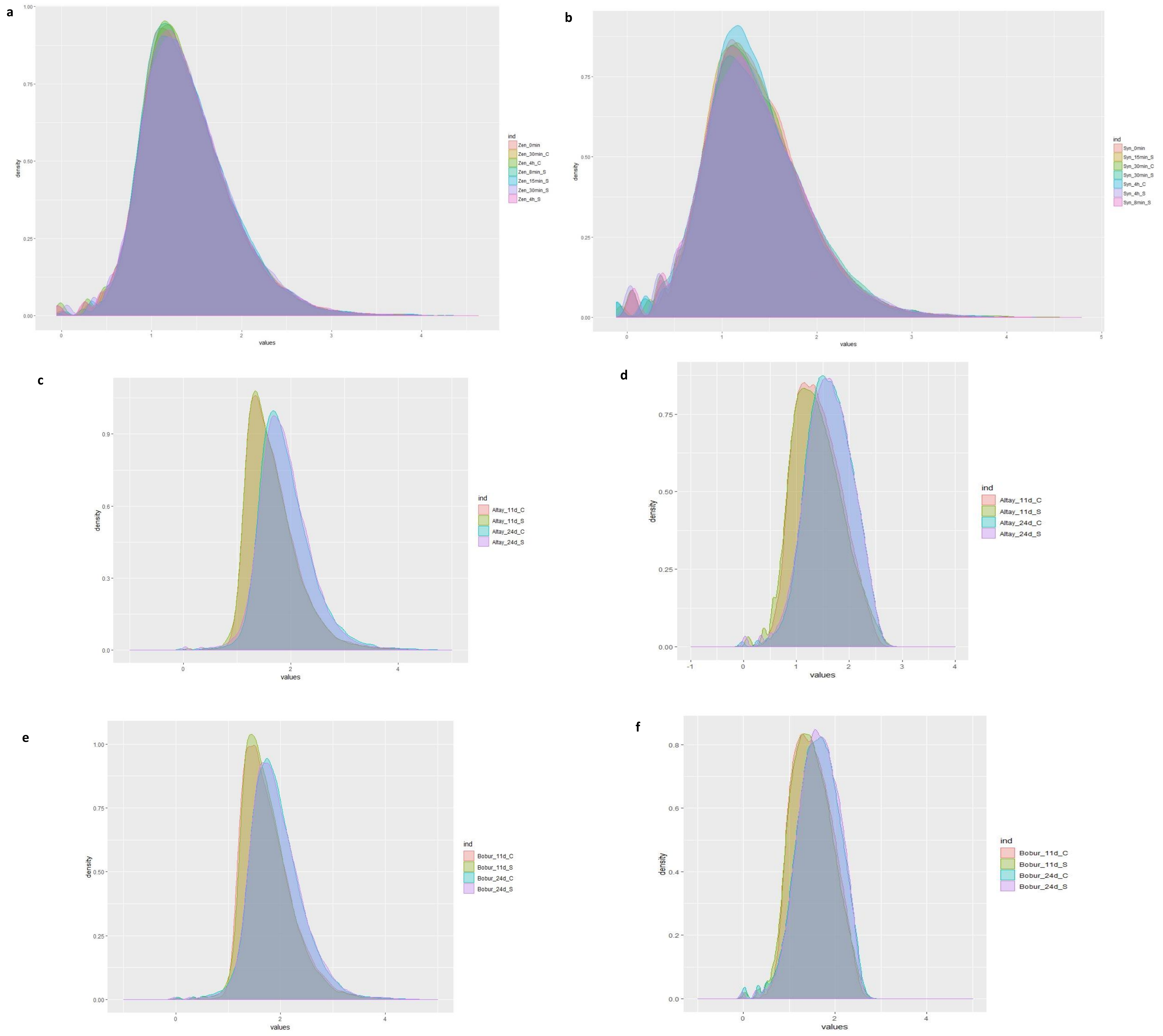


**Figure S1**. Density plots with the $\log_{10}$ normalized expression values of the libraries from the four genotypes studied. a) Zentos, b) Syn86, c) Altay2000 without deduplication, d) Altay2000 deduplicated, e) Bobur without deduplication and f) Bobur deduplicated. C= Control library, S= Stress library.
